# Impact of Blended NPSB Fertilizer Rates on Sugarcane (*Saccharum officinarum* L.) Varieties: Seed Cane Yield and Quality Insights from North Western Ethiopia

**DOI:** 10.1101/2024.08.02.606441

**Authors:** Solomon Ali, Bahiru Belay, Mesfin Abate

**Author notes:** Corresponding author: Solomon Ali Gebeyehu;.

## Abstract

Sugarcane primarily cultivated for sugar production and other multiple uses. A field experiment was conducted at Tana Beles sugar project, North western Ethiopia during 2021 and 2022 cropping season to determine the optimum rate of NPSB fertilizer rate on three sugarcane varieties. The treatments were laid out in factorial randomized complete block design arranged with three replications. The experiment was arranged with five levels of NPSB blended fertilizer (0, 200, 260, 320 and 380 kg ha^-1^) combined with three sugarcane varieties (NCO-334, N-14 and C86/56). Among the parameters of seed cane crop; germination percent stalk weight, stalk diameter, node length, inter node number, plant height, stalk population, and sett yield for growth and yield parameters and sett moisture content, sett nitrogen content, reducing sugar content and total sugar content for seed cane quality parameters significantly affected by applied NPSB fertilizer, varieties and their interaction (p<0.05). Brix% and pol% were not significantly affected by different rates of NPSB fertilizer rates and varieties (p<0.05). The highest leaf area index, plant population, sett yield, average cane weight, seed cane moisture content and total nitrogen content was attained with 380 kg ha^-1^NPSB fertilizer applied on variety NCO-334 and N-14. Maximum population stand, average plant height, and sett yield of verity C86/56 were recorded at 320 kg ha^-1^ NPSB fertilizer level. Sett yield, were positively correlated with germination percent, population stand count, inter node number, seed cane weight, seed cane diameter reducing sugar content, total sugar yield and sett moisture content. Therefore, it is advisable to recommend 380 kg NPSB ha^1^ for variety NCO-334 and N-14, and 320 kg NPSB ha^1^ for variety C86/56 with application of 160 kg ha^-1^ urea at the age of two month and half for effective seed cane production. It was aimed to fill the seed cane fertilizer rate problems of different sugarcane varieties.

## 1. INTRODUCTION

Sugarcane (*Saccharum officinarum* L.) belongs to the family *Poaceae* is one of the most important cash crop grown extensively all over the world covering 26 million hectares and 1.9 billion metric ton annual sugar production in more than 110 countries[1]. It is a monocotyledonous, tall-growing perennial tropical grass that tillers at the base to produce un- branched stems growing up to 4m. It is one of the most efficient converters of solar energy into sugar and other renewable forms of energy and hence produced primarily for its ability to store high concentrations of sugar in the internodes of the stem [2].

Sugarcane is primarily a tropical plant which is able to grow between 22°N and 22°S, and some up to 33°N and 33°S [3]. In terms of altitude, sugarcane crops are found up to 1,600 m.a.s.l close to the equator in countries such as Colombia, Ecuador, and Peru[4]. It usually requires between 8 to 24 months to reach maturity and temperatures high enough to permit rapid growth for 8 or more months depending on location [5] [6].

About 80% of the sugar produced globally comes from a species of sugarcane called *S.officinarum* and hybrids using this species. Sugarcane accounts for 79% of sugar produced; most of the rest is made from sugar beets. While sugarcane predominantly grows in tropical and subtropical regions, sugar beets typically grow in colder temperate regions. The average yield of cane stalk is 60–70 tonnes per hectare (24–28 long ton/acre; 27–31 short ton/acre) per year. However, this figure can vary between 30 and 180 tons per hectare depending on the knowledge and crop management approach used in sugarcane cultivation[6].

Seed cane production, which is an integral component of sugar production, often receives less priority than the commercial crop plants in many sugar cane plantations[7]. Most of the research works on seed cane has focused also on the mechanics of cutting and fungicidal treatments of setts. Therefore, little effort has been made to improve cultivation of seed cane[8] [9]. In order to maintain a uniform stand of sugar cane that ultimately produces high cane and sugar yield [10] [11].

Most research works focus on NP requirements of crops, limited information is available on various sources of fertilizers K, S, Zn and B and other micronutrients. Therefore, application of other sources of nutrients beyond Urea and Di-ammonium Phosphate (DAP), especially those containing K, S, Zn and other micro-nutrients could increase sugarcane productivity [12]. This can be achieved by application of blended fertilizers, the mechanical mixture of two or more granular fertilizer materials containing N, P, K and other essential plant nutrients such as S, Zn, and B, recently known to Ethiopia. [13].

Sundara reported that sugarcane makes heavy demand for plant nutrients. An average of 1.0 kg N, 0.6 kg P2O5 and 2.25 kg K2O are removed by a tone of sugarcane [14]. According to Ambachew and Tadesse, 1.20 kg N/ha and 0.80 kg P2O5/ha is required to produce an expected cane yield of 1 ton ha^-1^ In Finchaa Sugar Estate, 114 kg/ha and 115 kg P2O5 ha^-1^ is applied for plant cane to supplement the required nutrients [15].

Beles Sugar Development Project is new to sugarcane cultivation and had no site-specific fertilizer recommendations developed with respect to sugarcane varieties. In this regard, tentative fertilizer recommendation (250 kg DAPha^-1^ and 185 kg Urea ha^-1^) were recommended to Beles Sugar Development Project based on experiences of Finchaa Sugar Estate and limited number of soil samples [8] [13]. In 2016 the project research team prepared the first standard operation manual including the fertilizer (stating as 320 kg ha^-1^ NPSB and 160 kg ha^-1^ urea) as a state recommendation simply calculating the nitrogen and phosphorus percentage without considering the effect of sulphur and boron[16].

This study addresses limited knowledge on soil fertility especially concerning blended fertilizer usage specifically to different sugarcane varieties in the study area for increasing seed cane productivity. Tana Beles sugar development project currently uses similar blanket recommendations for seed cane, commercial, and ratoon cane production without regard to variety and soil characteristics. As a new developing sugar factory, use of scientific recommended blended fertilizer rate has paramount economic advantage to produce a healthy, vigorous and required quantity and quality seed cane for commercial cane production. For sugarcane particular seed cane crop; the economic optimum rate of NPSB is not well known in the study area. Therefore; this research fill the information gap on the blended fertilizer rate for seed cane production at TBSP.

### Objective

- To investigate the effects of blended (NPSB) fertilizer rates for set yield, yield and quality traits of seed cane
- To investigate the interaction effects of varieties and different rate of NPSB fertilizer on yield and yield components of seed cane
- To determine economic optimum rates of NPSB fertilizers for the productivity of seed cane

## 2. MATERIALS AND METHODS

### 2.1 Site Description

The study was conducted at Beles Sugar Development Project, which is found in an altitude of 1119 m.a.s.l in Amhara and some part of Benshangul Gumuz regional states of Ethiopia. The experiment was conducted in 2021 and 2022 main cropping season The average annual rainfall of 1490 mm. The minimum and maximum temperature of the area is between 16.4 and 32.5°C, respectively. The research area constitutes a vertie soil having 44.4% clay, 31.2% silt and 24.4% sand. It was carried out with a dragline sprinkler irrigation system.

### 2.2 Treatment and Experimental Design and Procedure

The treatment consisted of five rates of NPSB fertilizer levels (0, 200, 260, 320, and 380 kg ha^-1^) and three cane varieties namely NCo_334, N-14 and C86/56

**Table 1.** Description of varieties’.

| Variety | Breeder<br>(Country) | Brunching<br>nature | maturity type | Seed rate<br>(qt/ha) |
| --- | --- | --- | --- | --- |
| NCO-334 | Combitor/India | Highly<br>brunching | intermediate | 90 -95 |
| N-14 | South Africa | Moderate | intermediate | 95 -100 |
| C86/56 | Cuba | Low | early | 100- 120 |

Treatments are arranged factorially and laid in a randomized complete block design with three replications. Nitrogen was applied with the recommended rate (134kg ha^-1^).

### 2.4 Soil Physico-chemical analysis

Composite soil samples for the study area were collected based on the standard and was subjected to physico-chemical analysis to evaluate the texture, structure, organic carbon, organic matter, pH, CEC, and macro nutrient content including nitrogen and phosphorus before planting and after harvesting according to the standard operation of the project prepared in 2016

### 2.5 Data collected

The following data were collected germination percentage, stalk length, cane stalk diameter/girth, number of internodes per seed cane stalk, seed cane weight per stalk, seed cane stalk population/tillers, seed cane node length, set yield per hectare, total nitrogen content, sugar analysis, cane moisture content and total soluble solids (TSS)

### 2.6 Statistical Analysis

The data was subjected to the General Linear Models Procedure (GLM) using SAS Version 9.4 software statistical package (SAS, 2019) following a procedure appropriate to the design of the experiment. The treatment means was separated by using the least significant difference at 5% significance.

### 2.7. Partial budget /Economic analysis

An economic analysis was done using partial budget procedure described by CIMMYT, The cost of NPSB and seed cane was considered during planting. The net returns (benefit) and other economic analysis was done based on the formula developed by [17].

## 3. RESULT AND DISCUSSION

### 3.1 Soil physical and chemical Analysis

Soil analysis was made for the sample collected from the research site before fertilizer application from the experimental site and after harvesting of the seed cane for the determination of the major soil chemical properties at Pawe Soil Research and Laboratory center. The soil of the study site is acidic in reaction but low in exchangeable acidity. Moreover, the total nitrogen and available phosphorus of the soil were low. Therefore, the result revealed that soil of the experimental site is deficient in plant nutrients. The pH of the soil is medium acidic before applying the treatments but during harvesting the pH, organic matter, organic carbon and cation exchange capacity slightly drops with increasing the level of NPSB fertilizer. While available nitrogen, phosphorus, sulphur and boron level increased with increased level of NPSB fertilizer.

### 3.2. Effects of NPSB Fertilizer on Growth and Yield Parameters of Seed Cane varieties

#### 3.2.1 Germination Percentage

The highest germination percentage (75.7%) was recorded in variety NCO-334 treated with 260 and followed 320 kg ha^-1^ of NPSB fertilizer whereas the lowest percent of germination was recorded (53.2%) in variety C-86/56 without NPSB fertilizer treatment. It shows 22.5% difference from the highest to the lowest percentage on the sprouting ability of seed sets (**Table 2**). This was mainly due to difference in bud nature of which was genetical between the varieties. The Present finding is in harmony with [18]. [19], also reported that the nutritional status of cane stalk/sett had marked influence on germination of sett for the subsequent commercial crop as it is afforded the energy required for sprouting of bud and young shoot till it was established on its own. Wubale [8] also revealed that, use of N fertilized planting material for commercial cane production showed significant difference (p<0.01) in sprouting ability of the cane in Luvisol (light soil) of Tana Beles. Similar result was obtained by Sime at Finchaa sugar estate [20].

#### 3.1.2 Inter Node Length and Inter Node Number

The analysis of variance value indicates that there was significant difference between the interaction and levels of NPSB fertilizer (p<0.05). However varietal difference did not show significant difference. The highest node length (12.19) was recorded at variety C86/56 treated with 260 kg ha^-1^ NPSB level of the trial which is significantly different from all treatment combinations except variety NCO-334 and N-14 treated with 320 and 380 kg ha^-1^ NPSB fertilizer and variety C86/56 treated with 320 kg ha^-1^ NPSB fertilizer (**Table 2**). Whereas, the lowest inter node length (9.21) was recorded at the combination of NCO-334 without the application of NPSB with 2.98 cm shorter than the longer internode which was significantly different from variety N-14 and C86/56 receiving 260, 320 and 380 kg ha^-1^ NPSB fertilizer. The difference was due to increase in the level of the fertilizer which increases early vegetative growth which again increase inter node length and number.

The analysis of variance showed that there was significant difference between the interaction (p<0.05). The highest number of inter node (14.43) were recorded on variety C86/56 treated with 260 kg ha^-1^ NPSB fertilizer which was significantly par with variety C86/56 received 0, 200, 320, 380 kg ha^-1^ and variety NCO-334 treated with 0, 320 and 380 kg ha^-1^ NPSB fertilizer while the rest treatment combinations were significantly different (**Table- 1**). On the other hand the lowest inter node number (11.36) were recorded at variety N-14 without NPSB fertilizer which was significantly different to all treatment combinations except application of 200, 260, 320 and 380 kg ha^-1^ of NPSB fertilizer on variety N-14 and application of 200 and 260 kg ha^-1^ NPSB fertilizer on NCO-334. It was evident that from table 1 there was three inter node number difference for each single cane with the highest to lowest number of internodes.

This was in harmony to the work of Dereje *et al*. [12] who reported that blended fertilizer effect with different rates on ratoon cane production. From this result it was observed that increasing blended fertilizer rate up to same extent increases the number of nodes and length of internodes which have an advantage to get longer stalk which will resulted in high cane yield. The present finding is in line with [21] that application of Zn and B increased the average length, girth, internodes number, and weight cane per stalk, number of millable canes and yield of sugarcane. From this result it was observed that blended fertilizer increases the number of internodes which have an advantage to get longer stalk which will resulted in high cane yield, biomass yield and sugar recovery.

#### 3.1.3. Plant Height

The main effect of NPSB fertilizer rate and Varieties as well as interaction effect significantly (P < 0.01) affected plant height (Appendix table 2). The highest increment in height was observed on C86/56 seed cane plants receiving 320 kg ha^-1^ of NPSB fertilizer followed by variety C86/56 receiving 260 kg ha^-1^ NPSB fertilizer. While the shortest plant height was observed from a variety N-14 treated with 200 kg ha^-1^ of NPSB fertilizer (1.62 m) (**Table 2**). The increased in the plant height due to NPSB fertilizer was caused by increase in number of nodes or inter nodes elongation or both. The highest seed cane height was observed on Variety C86/56 which is early maturing ones whereas the lowest height record was observed on late maturing ones (N-14). Variety C86/56 receiving 320 kg ha^-1^ of NPSB fertilizer) was 27% greater in seed cane height from the control of the same variety and 36.4% greater than variety N-14 treated with 200kg ha^-1^ NPSB fertilizer which is the shortest value in height of the trial. Similar result was reported by Episten [22] on the increase in plant height with respect to increased NPSB application rate indicates maximum vegetative growth of the plants under higher levels of NPSB availability. More over the smallest plant height was achieved from untreated plant[23]. This result was also supported by the finding of [12] as the application of N at early growth stage of the seed cane plants enhanced better vegetative growth. This result was also in agreement with the report of [24] [16] who reported that plants deficient in N exhibited retarded growth. Similarly, Sime [20] reported a close relationship between growth and applied nitrogen in which high amount of applied N resulted the highest plant height and vice versa on the experiment done at Finchaa. In addition, a study done on the effect of nitrogen and phosphorus fertilizers on growth and yield of quality protein maize at India by [11] revealed that, the growth parameters like plant height, leaf area index and dry matter production was significantly affected by the application at different levels of fertilizers [11]. According to their finding, maize crop fertilized with high amount of NPSB fertilizer had significantly resulted in long statured plants compared to lower nitrogen levels including untreated check.

**Table 2.**
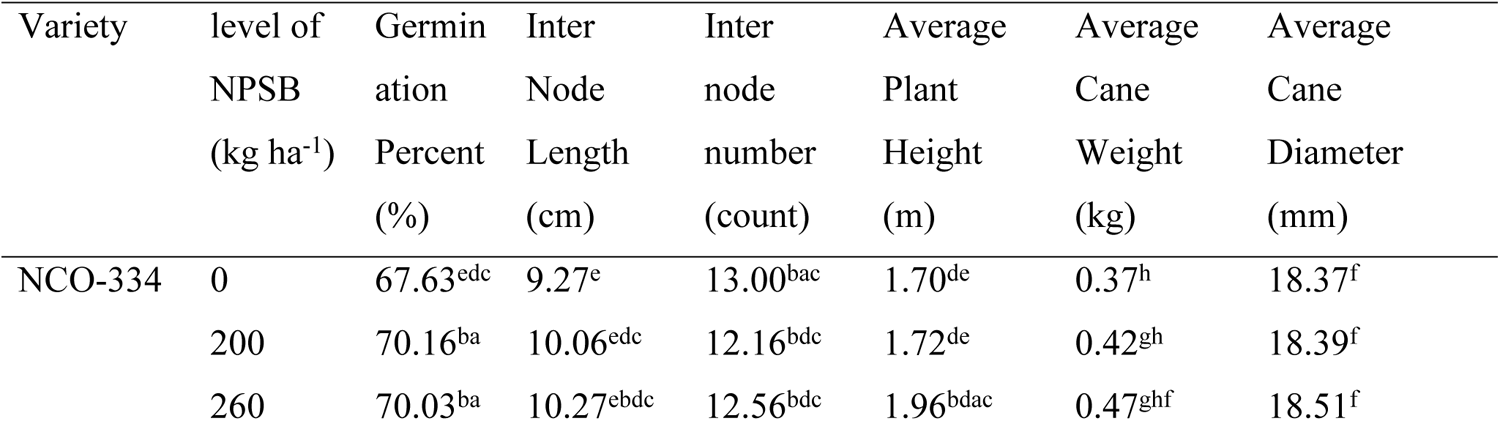
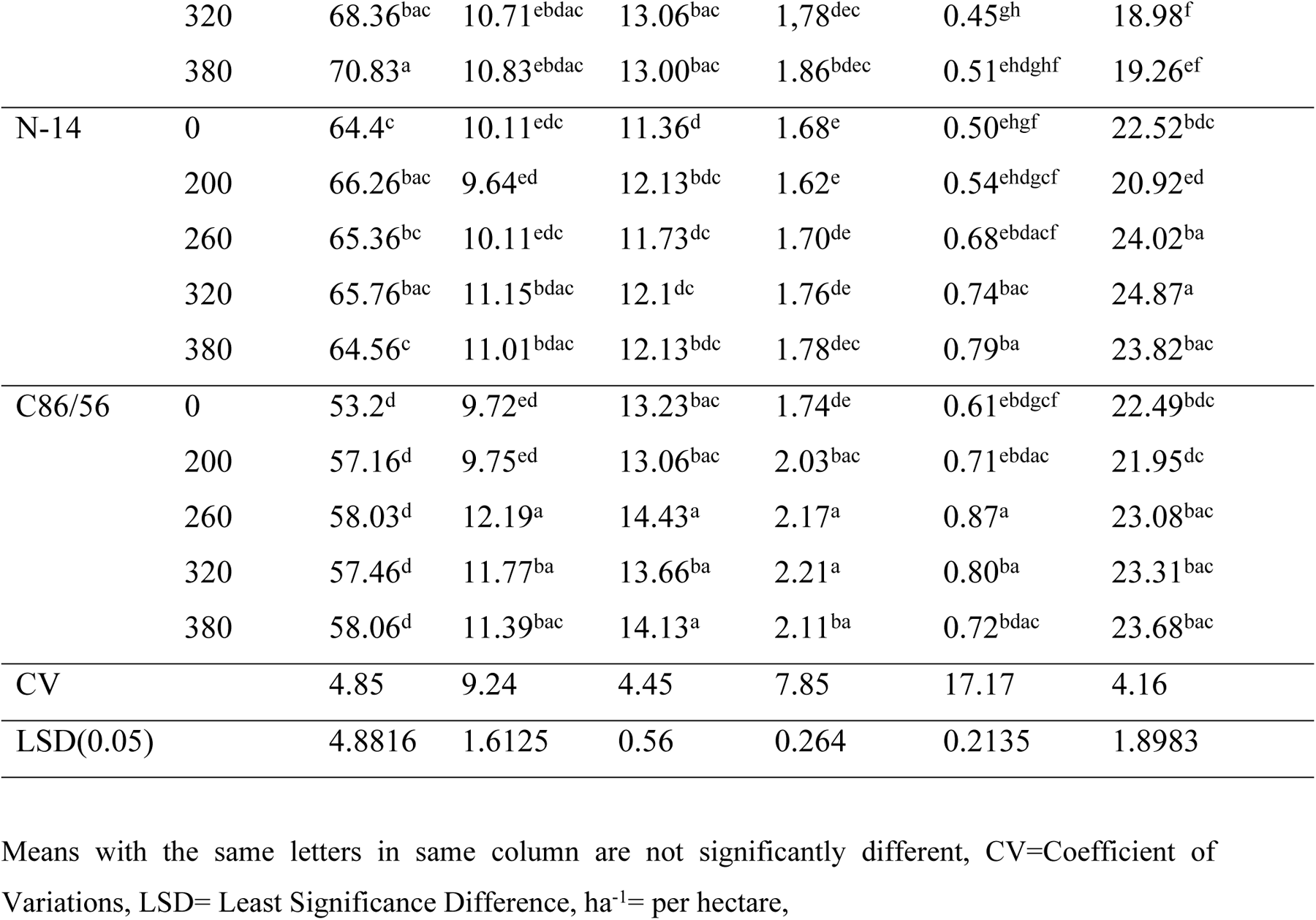
Interaction effect of varieties and NPSB fertilizer on Inter node length, Inter node number, plant height, cane weight and average cane diameter.

#### 3.1.4 Cane Weight per Stalk and Diameter

The analysis of variance showed that the effect due to NPSB blended fertilizer rate on weight per stalk were significant (P<0.05) and the interaction effect of NPSB and varietal effect was highly significant (P<0.01) on weight per stalk.

The highest average weight (0.51 kg) was recorded, by treating the seed cane variety NCO-334 treated with 380 kg ha^-1^ of NPSB but this was statistically the same with that of treated by 200, 260, and 320 kg of NPSB ha^-1^ for variety NCO-334, 0.79 kg for treatment combination of N- 14 with 380kgs of NPSB fertilizer which was significantly different from the control and treatment six : which receives 200kg of NPSB for N-14 and 0.87kg ha^-1^ for C86/56 which is significantly different from variety C86/56 without NPSB and variety N-14 receiving 200 kg of NPSB fertilizer get the maximum cane weight with the highest level of nitrogen and NPSB fertilizer level respectively [8].

The analysis of variance showed that seed cane girth and seed cane weight were highly significantly affected (P<0.01) by the application of NPSB blended fertilizer rate, variety and the interaction (**Table 2**). The highest stalk diameter (24.87mm) was recorded under application of 320 kg ha^-1^ NPSB blended fertilizer on variety C86/56. The least seed cane diameter was recorded at control treatment (0 kg ha^-1^ NPSB fertilizer for variety NCO-334). However, it was statistically at equivalence for all of the NPSB blended fertilizer treatment results of variety NCO-334 and significantly different from both N-14 and C86/56 for all fertilizer rate of applications of the experiment (**Table 2**). Moreover, high plant population on variety NCO-334 produced thinner cane stalks due to crowding effect, whereas low plant population on variety C86/56 produced thicker cane stalks because of the availability of wider space. This finding is in agreement with the works of [5] in which the wider spacing’s there was a higher stalk weight than in the narrower spacing’s. Moreover, Jiang [25] also stated that as density of planting increases stalk weight decreases.

Humbart [24] observed that the cane length and diameter, number of tillers per plant, cane yield and sugar recovery increased with the application of nitrogenous fertilizers in the sugarcane varieties. Mohanty [26] also described that, nitrogen and phosphorus containing fertilizer is the key nutrient element influencing sugarcane yield and quality. It is required more for vegetative growth, i.e. tillering, foliage formation, stalk formation, stalk growth (internodes formation, internodes elongation, increase in stalk girth and weight) and root growth.

Furthermore, the stalk girth plays an important and dominant role in improving cane yield per unit area, which could be due to the indirect increase in stalk weight [27].

#### 3.1.5 Number of population and Leaf area index

The analysis of variance showed that all varieties and interactions differed significantly (p<0.05) among each other for number of populations. The maximum number of populations count ware rescored on 320 kg ha^-1^ of NPSB fertilizer applied for variety NCO-334 and it was statistically at par with variety NCO-334 treated with 200 and 260 kg ha^-1^ NPSB fertilizer. On the other hand, the minimum numbers of population were noticed on variety C86/56 having no fertilizer application which was statistically at par with all treatments except variety N-14 treated with 200 kg ha^-1^ NPSB fertilizer, variety C86/56 treated with 200 kg ha^-1^ and 260 kg ha^-1^ NPSB fertilizer. The present study is consistence with the work conducted by Majeedano [28] reveals similar results on number of millable canes. . The presence of varietal differences in number of millable canes was reported by [29]. Similar to this result, Tadese [18] also observed variation among varieties on number of population canes. One of the main factors affecting tillering of seed cane are inorganic fertilizer applied, even though the number of developing tillers per stool varies with variety and the growing conditions. Moreover they also revealed that increment on the amount of NPSB blended fertilizer increases early tillering and good stand of seed cane growth which is the source of population number [29] [13].

In relation to treatment effects on leaf area index there was significant difference between varieties, between NPSB levels and their interaction (p <0.01). The highest leaf area index was recorded at variety C86/56 receiving 380 kg of NPSB blended fertilizer recording a value of 5.41 which is significantly different from all NPSB fertilizer levels applied on variety N-14 and variety NCO-334 without NPSB fertilizer (control) (**Table 2**). Whereas variety N-14 receiving 200 kg of NPSB blended fertilizer was the lowest value (2.85) in the trial which was significantly as par with variety N-14 treated with 200, 260, 320, and 380kg ha^-1^ NPSB fertilizer levels and variety NCO-334 without the application of NPSB fertilizer. Increasing the levels of NPSB fertilizer level up to 380 kg ha^-1^ increases the value of leaf area index which was due to good vegetative growth stand of the cane varieties receiving the indicated level of the fertilizer [30].

**Table 3.**
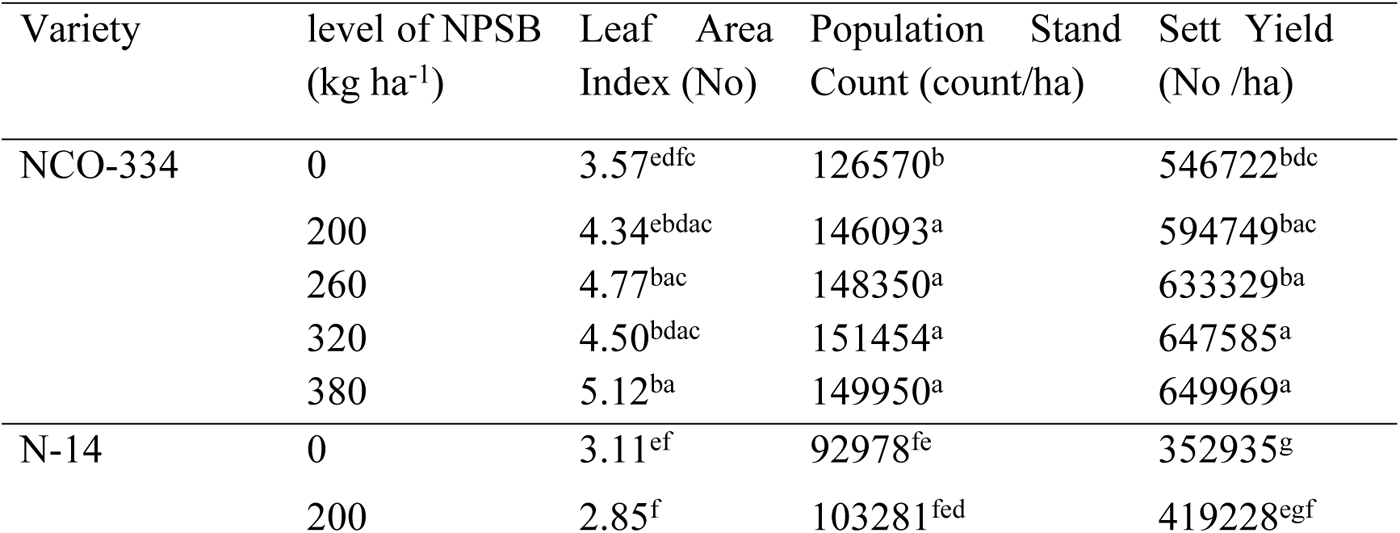
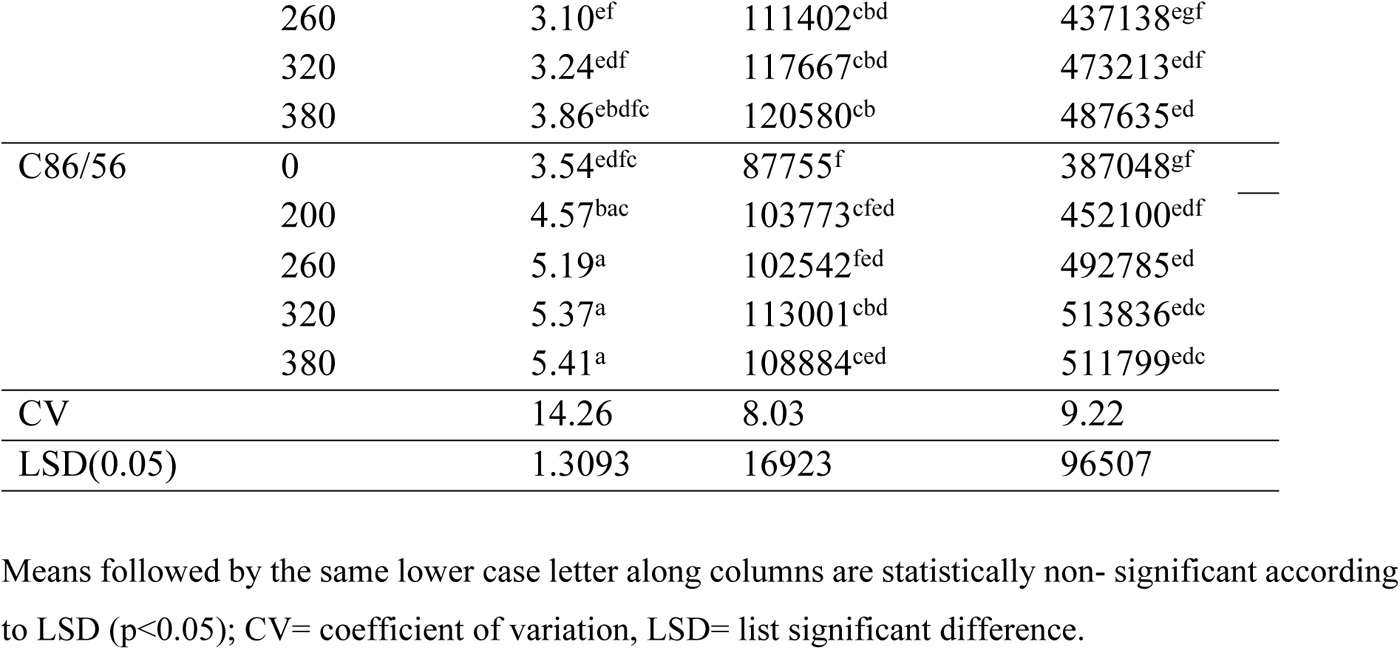
Interaction effect of varieties and NPSB fertilizer on leaf area index, population number and sett yield.

#### 3.1.6 Sett Yield

The response of sett yield of seed cane showed significant (p < 0.01) difference for different rates of NPSB fertilizer, variety and interaction effects. The highest sett yield (649,969) in number each having three bud had recorded at variety NCO-334 treated with NPSB fertilizer of 380 kg ha^-1^ which was similar to population number. It is significantly different from all treatments except a combination of variety NCO-334 with 200, 260, and 320 kg ha^-1^ of NPSB fertilizer levels. While the lowest number of seed cane yield (352,935) was recorded at the variety N-14 without NPSB fertilizer application which was statistically the same (p<0.05) with variety N-14 treated with 200 and 260 kg ha^-1^ and variety C86/56 without NPSB fertilizer.

This result was similar with the work of Sime [20] for sett yield in which significant differences between treatments for sett yield was observed on experiment conducted at Finchaa on different rate and time of N application. The same result was also obtained from the work of Zeleke [13] and Hussain [31] for seed cane yield. As they reported from the experiments conducted at Finchaa and Wonji-Shoa. Zeleke [13] showed that yield increment of 26 ton ha^-1^ with increasing the level of nitrogen and phosphorus fertilizer from 90 to 136 kg ha^-1^. While Abdurrahman [32] Revealed that Application of 150kg ha^-1^ ammonium Sulphate/feddan (36 kg ha^-1^ sulfur) increased sugarcane yield and showed significant difference. Demmsie [33] also reported that blended fertilizer treatment with the rate of (250kg ha^-1^ blended + 94kg N) ha^-1^ at one month after harvest resulted in higher ratoon cane weight per stalk, stalk girth, cane yield, sugar yield, node length, stalk population and node number on sugarcane.

### 3.2 Seed Cane Quality Parameters

#### 3.2.1 Moisture Content

Seed cane moisture content of the was highly significantly (p<0.01) influenced by both varietals, NPSB level and their interaction. The highest moisture content of the sett (79.44) was observed for variety N-14 treated with 380 kg ha^-1^ NPSB fertilizer rate, which was statistically at par with variety NCO-334 treated with 320 and 380 kg ha^-1^ of NPSB fertilizer level and significantly different from other treatments combinations. Whereas the lowest percent of seed cane moisture content (77.1%) was recorded at variety C86/56 treated without NPSB fertilizer level which was statistically the same with NCO-334 treated without, and 200 kg ha^-1^ NPSB fertilizer and variety C86/56 receiving 200 and 260 kg ha^-1^ of NPSB fertilizer level. The trial showed that the higher percentage of moisture content in seed cane is a good quality indicator. It implies that application of the project recommended dose and above gives higher moisture content for all varieties taken for the trial. The higher moisture and nitrogen content of seed cane was recorded at the higher the application of NPSB fertilizer dose to the varieties taken under the experiment while the lower moisture, nitrogen and reducing sugar content were recorded to the control.

The present finding is consistent with [26, 27] reported that additional fertilizers given to sugarcane crop planted exclusively for seed purpose improved seed quality by enhancing sett nutrient status and sett moisture content. Vilela [2] also reported that sets with higher moisture content give quicker and higher germination and the seedlings emerging from such setts establish quickly and grow vigorously. Singh [6] also reported that optimum level of sett moisture content for rapid germination was 72 to74%. In addition, Srivastava [34] [35]also described standards of seed cane moisture content and suggested that it should not be less than 65% on weight basis. In general, the average stalk/sett moisture content of all treatments in the study was greater than that of the critical sett moisture content (50.3%) for germination of buds on seed cane set observed in **Table 6**.

#### 3.2.2 Seed Cane TSS, POL% and Total Sugar Content

From the analysis of variance, only varietal difference was significant (p<0.05) for seed cane total soluble solid and percent of polarity whereas NPSB fertilizer and interaction did not have influence for total soluble solid. However, for total sugar content a significant difference was observed between varieties, NPSB blended fertilizer levels and their interaction (p<0.01).

Comparing the varieties on which the trial was conducted, C86/56 showed the highest pol and ^0^ brix recording 13.641 and 15.66 respectively. While the lowest percent of polarity and ^0^ brix was observed on Varity N-14 recording 11.53 and 14.3 respectively. However intermediate ^0^ brix and percent of pol accumulation was observed on variety NCO-334 with significantly differing from N-14 (**Table 3**).

This finding is in close confirmation with the findings of [36] [37]. These results are also in accord with those of Sarwar [7] who reported that lower rate application of NPSB fertilizer resulted in poor juice quality (low brix and pol value) in matured cane but in contrary to this, it is a good quality for seed cane plants in maintaining food reserve for the germinating buds[38]. According to [39] ^0^brix and pol percent are genetic character hence varietal difference was significantly different from each other.

Ibrahim [40] who described an inverse relation between the increasing solid fertilizer and decreasing pol% in juice. [14] evaluated the Pol of several sugarcane cultivars over the 2009/2010 cropping season and found a variation in apparent sucrose content in all cultivars, characterized by an increase with the advancement of phenological stages. According to Santos [41], when it comes to mature sugarcane, there is a close relationship between apparent percentage of soluble solids and sucrose content in the solution and sugarcane is considered mature with minimum of 18° Brix, among other factors.

**Table 4.** Main effects of varieties on seed cane total soluble solid (^0^Brix) and Pol%.

| Treatments | TSS° brix | Pol% |
| --- | --- | --- |
| Varities |  |  |
| NCO-334 | 15.6<sup>a</sup> | 12.946<sup>a</sup> |
| N-14 | 14.3<sup>b</sup> | 11.53<sup>b</sup> |
| C86/56 | 15.66<sup>a</sup> | 13.641<sup>a</sup> |
| LSD(0.05) | 0.693 | 0.71 |
| CV | 1.79 | 9.06 |

Means with the same letters in same column are not significantly different, CV=Coefficient of Variations, LSD= Least Significance Difference, ha<sup>-1</sup>= per hectare,

The result obtained in this study showed there was significant difference (p<0.05) for interaction. Whereas main effects did not have significance effect for recoverable sugar content. The highest value of recoverable sugar content (10.55 mg 100 g ^-1^) was recorded with variety NCO-334 treated with 380 kg ha-1 of NPSB fertilizer level which is statically different to variety N-14 and C86/56 treated to all levels of NPSB fertilizer rates of the trial. Whereas the lowest level of estimable/recoverable sugar content (9.25 mg 100 g ^-1^) of the cane was recorded at variety N-14 without NPSB fertilizer, which was statistically different from variety NCO-334 treated with the five levels of NPSB fertilizer (0, 200, 260, 320 and 380 kg ha^-1^). There was 1.3% of sugar yield difference between NCO-334 treated with 380 kg ha-1 and the lowest value recorded on variety N-14.

The present finding was consistent with the work of [8] [23] reported that blended fertilizer treatment with the rate of (250kg ha^-1^ blended + 94kg N) ha^-1^ at one month after harvest resulted in higher cane weight per stalk, stalk girth, cane yield, sugar yield, node length, stalk population and node number on sugarcane plant. The highest sugar yield was obtained at the rate of 136 kg N/ha and 138 kg P_2_O_5_/ha. At this rate, sugar yield increased by 29.14% over the conventional treatment (nearly 182 kg N/ha). The lowest sugar yield was obtained at sole application of 90 kg N/ha applied.

Negesse [3] revealed that the main effect and interaction effects of blended fertilizers at different levels of phosphorus were highly significantly (p<0.001) for parameters like sucrose yield but not for cane yield and Brix% [13].

#### 3.2.3 Reducing Sugar Content

The analysis of variance there was significant difference between interaction and main effect varieties (p<0.01), however application of different NPSB fertilizer levels did not show significant difference in the trial. The highest value 2.84 mg 100 g^-1^ was recorded at a variety N-14 treated with 260 kg ha^-1^ of NPSB blended fertilizer which is statistically different with variety NCO-334 treated without and 200 kg ha-1 NPSB fertilizer and variety C86/56 treated with all levels (0, 200, 260, 320 and 380 kg ha-1) of NPSB fertilizer (**Table 4**). Whereas the lowest value 1.87 mg 100 g^-1^ was recorded at variety C86/56 receiving 200kgha^-1^ of NPSB blended fertilizer which is statistically the same with variety C86/56 treated to different levels of NPSB fertilizer (**Table 5**).

The application of 260 kg ha^-1^ of NPSB blended fertilizer at planting and 225 kg/ha urea at top dressing may have favored vegetative growth, delayed maturation and reduced percentage of sucrose by increasing the content of reducing sugar [27].

#### 3.2.4 Seed Cane Nitrogen Content

Nitrogen content of seed cane of the trial was significantly (p<0.05) affected by different for both varietal, NPSB blended fertilizer as well as interaction effect. The highest nitrogen content of the trial was recorded with the application of 320 followed by and 380 kg ha^-1^ for varieties NCO-334 and N-14 and 380 kg ha^-1^ for C86/56 (**Table 5**) which was statistically different from each varieties treated with 0, 200 and 260kg ha^-1^ NPSB fertilizer levels. Whereas the lowest value was recorded for variety C86/56 without NPSB fertilizer application and which was statistically different from all treatment combinations except variety C86/56 treated with 200 kg ha^-1^.

The nutritional status including nitrogen content of cane stalk/sett had marked influence on germination of sett for the subsequent commercial crop as it is afforded the energy required for sprouting of bud and young shoot till it was established on its own [42]. Therefore, high nitrogen, starch and glucose contents are essential for good seed though their content varied with different varieties.[40]

**Table 5.**
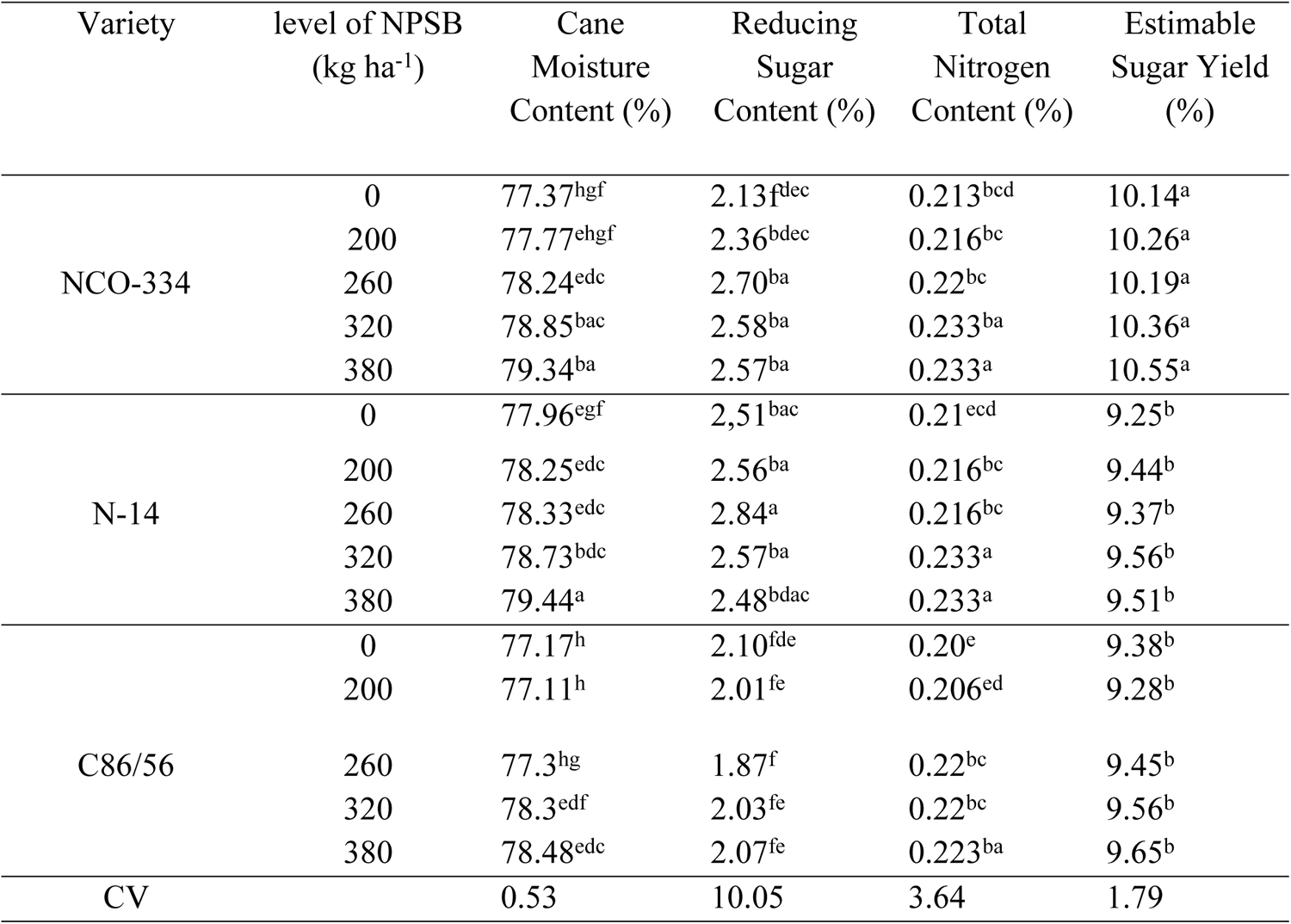
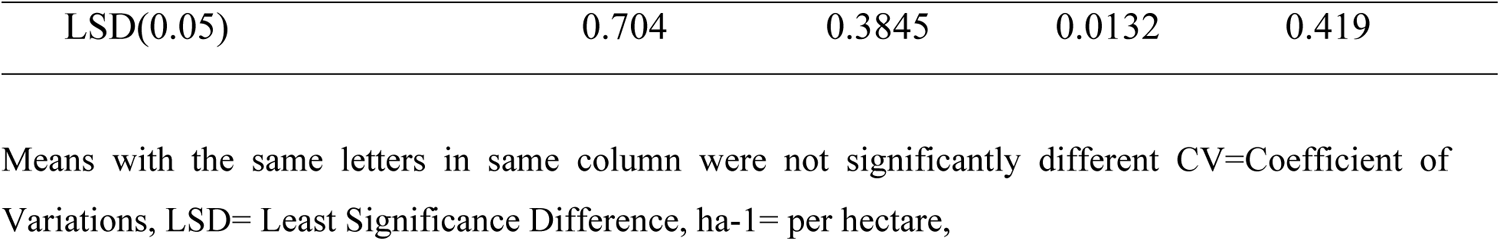
Interaction effect of varieties and NPSB fertilizer on seed cane moisture content, total soluble solid, TSS), pol%, reducing sugar content and total nitrogen content.

### 3.3 Partial budget/Economic Analysis

In this experiment, high values of net benefit ($7,518.89 ha-1) and BCR (8.28) were obtained at 380 kg NPSB ha-1 fertilizer rate. Increasing NPSB fertilizer rates from 0 to 380 kg/ha increases the net gain from $4,317.57 to $7,518.89, from $2,723.19 to $5,950.87, and from $2,909.11 to $6,067.54 for NCO-334, N-14, and C86/56 respectively. From **Table 5**, it was evident that it is possible to get 74.2% and 118.5% additional benefit when we treat 380 kg/ha NPSB fertilizer with 160 kg/ha of urea for seed cane variety NC0-334 and N-14 respectively. However, variety C86/56 scored the highest additional benefit (113.2%) when 320 kg/ha NPSB and 160 kg urea fertilizer were applied.

### 3.4 Correlation analysis

From **table 7** set yield was positively and significantly correlated with seed cane population number (r=0.907***), seed cane leaf area index (r=0.534***), seed cane diameter (r=0.4630**) seed cane inter node number (r=0.377*), sugar yield (r=0.501***) and seed cane nitrogen content (r=0.291***)This is in confirmation with the work of [16] [30], a research conducted on 12 sugarcane varieties in Pakistan. Wubale, [12] also get positive correlation between vegetative parameters and leaf nutrient contents of seed cane. Birhanie [3] also showed that there were positive association between pol, brix% and sugar content of sugarcane plant for different levels and type of blended fertilizes.

**Table 6.** Correlation coefficients and the significant level between each parameter.

|  | GP | PSC | INL | INN | APH | ACW | ACD | LAI | SY | CMC | TSS | POL | RSC | ESY | TN |
| --- | --- | --- | --- | --- | --- | --- | --- | --- | --- | --- | --- | --- | --- | --- | --- |
| GP | 1 | 0.61851 | 0.09124 | 0.36552 | 0.37141 | 0.40715 | 0.51958 | 0.17766 | 0.42252 | 0.45419 | 0.01831 | 0.12647 | 0.52083 | 0.501 | 0.291 |
|  |  | <.0001 | 0.5511 | 0.0135 | 0.012 | 0.0055 | 0.0003 | 0.243 | 0.0038 | 0.0017 | 0.905 | 0.4078 | 0.0002 | 0.0004 | 0.052 |
| PSC |  | 1 | -0.00159 | 0.03899 | 0.01793 | 0.39134 | 0.56868 | 0.32676 | 0.90783 | 0.45035 | 0.13384 | 0.05921 | 0.32858 | 0.8008 | 0.845 |
|  |  |  | 0.9917 | 0.7993 | 0.907 | 0.0079 | <.0001 | 0.0285 | <.0001 | 0.0019 | 0.3808 | 0.6992 | 0.0275 | <.0001 | 0.001 |
| INL |  |  | 1 | 0.39901 | 0.64305 | 0.64851 | 0.3992 | 0.40504 | 0.155 | 0.18526 | 0.24459 | 0.27281 | 0.1608 | 0.0987 | 0.2793 |
|  |  |  |  | 0.0066 | <.0001 | <.0001 | 0.0066 | 0.0058 | 0.3093 | 0.2231 | 0.1054 | 0.0698 | 0.2913 | 0.5203 | 0.0628 |
| INN |  |  |  | 1 | 0.64501 | 0.47529 | 0.15588 | 0.58746 | 0.37792 | 0.13538 | 0.33605 | 0.45609 | 0.50076 | 0.1966 | 0.0038 |
|  |  |  |  |  | <.0001 | 0.001 | 0.3065 | <.0001 | 0.0105 | 0.3752 | 0.024 | 0.0016 | 0.0005 | 0.1954 | 0.9809 |
| APH |  |  |  |  | 1 | 0.60476 | 0.28325 | 0.67888 | 0.2409 | 0.12002 | 0.16668 | 0.2562 | 0.35568 | 0.0464 | 0.0623 |
|  |  |  |  |  |  | <.0001 | 0.0594 | <.0001 | 0.1109 | 0.4323 | 0.2738 | 0.0894 | 0.0165 | 0.7639 | 0.6846 |
| ACW |  |  |  |  |  | 1 | 0.77194 | 0.20193 | 0.17756 | 0.09896 | 0.02801 | 0.08063 | 0.26729 | 0.4781 | 0.0219 |
|  |  |  |  |  |  |  | <.0001 | 0.1834 | 0.2433 | 0.5178 | 0.8551 | 0.5985 | 0.0759 | 0.0009 | 0.886 |
| ACD |  |  |  |  |  |  | 1 | 0.019 | 0.46326 | 0.1169 | 0.3057 | 0.19313 | 0.14068 | 0.6358 | 0.0495 |
|  |  |  |  |  |  |  |  | 0.9014 | 0.0014 | 0.4444 | 0.0411 | 0.2037 | 0.3567 | <.0001 | 0.749 |
| LAI |  |  |  |  |  |  |  | 1 | 0.53412 | 0.05279 | 0.34268 | 0.40625 | 0.29604 | 0.3633 | 0.1613 |
|  |  |  |  |  |  |  |  |  | 0.0002 | 0.7305 | 0.0212 | 0.0056 | 0.0483 | 0.0141 | 0.2879 |
| SY |  |  |  |  |  |  |  |  | 1 | 0.36248 | 0.26405 | 0.24452 | 0.09842 | 0.8302 | 0.5064 |
|  |  |  |  |  |  |  |  |  |  | 0.0144 | 0.0797 | 0.1055 | 0.5201 | <.0001 | 0.0004 |
| CMC |  |  |  |  |  |  |  |  |  | 1 | 0.13291 | 0.1658 | 0.51499 | 0.2903 | 0.706 |
|  |  |  |  |  |  |  |  |  |  |  | 0.3841 | 0.2764 | 0.0003 | 0.053 | <.0001 |
| TSS |  |  |  |  |  |  |  |  |  |  | 1 | 0.97285 | 0.51498 | 0.2443 | 0.0163 |
|  |  |  |  |  |  |  |  |  |  |  |  | <.0001 | 0.0003 | 0.1053 | 0.9163 |
| POL |  |  |  |  |  |  |  |  |  |  |  | 1 | 0.64746 | 0.18 | 0.016 |
|  |  |  |  |  |  |  |  |  |  |  |  |  | <.0001 | 0.2365 | 0.916 |
| RSC |  |  |  |  |  |  |  |  |  |  |  |  | 1 | 0.1598 | 0.2977 |
|  |  |  |  |  |  |  |  |  |  |  |  |  |  | 0.2941 | 0.047 |
| ESY |  |  |  |  |  |  |  |  |  |  |  |  |  | 1 | 0.4214 |
| TN |  |  |  |  |  |  |  |  |  |  |  |  |  |  | 0.004 |

## 4. CONCLUSION AND RECOMMENDATION

The result of the study had a marked effect and showed a highly significant difference (p<0.05) for germination percent, seed cane weight, seed cane diameter, node length, inter node number, plant height, stalk population, leaf area index and sett yield for growth and yield parameters and sett moisture content, sett nitrogen content, total sugar content, reducing sugar content for seed cane quality parameters of the different NPSB blended fertilizer rates for the study area. Application of NPSB blended fertilizer along with of UREA fertilizer remarkably increased seed cane yield of sugarcane. From the fifteen (15) agronomic parameters measured in the experiment three of them (LAI, seed cane moisture content and total nitrogen content) showed highest record result at treatment 380 kg ha^-1^ NPSB fertilizer rate for the three seed cane varieties which is 60 kg NPSB fertilizer above from the projects standard.

The highest sett yield was recorded with application rate of 380kg ha^-1^ of NPSB and 160 kg ha^-1^ of urea fertilizer for variety NCO-334 and N-14, and 320 kg ha^-1^ NPSB and 160 kg ha^-1^ urea fertilizer for variety C86/56.

When we see the correlation, germination percent, population stand count, inter node number, seed cane weight, seed cane diameter, sett yield, reducing sugar content, total sugar yield and sett moisture content positively and strongly correlate each other.

The economic analysis of the trial indicates that application of 380 kg of NPSB at planting and application of 160 kg of urea gave the highest marginal rate of return (936%) and a net field benefit of $7,618.85, which is $287.92 greater than the project’s recommendation for variety NCO-334. Variety N-14 at the 380 kg NPSB fertilizer level gave a net field benefit of $6,045.98 and a 714% marginal rate of return, which is $527.62 greater than the estate recommendation prepared by the project research team. Whereas variety C86/56 gave the highest net field benefit of $6,295.77 and a marginal rate of return of 793% at the application rate of 320 kg/ha of NPSB and 160 kg/ha UREA, which is on par with the estate recommendation.

Accordingly, based on the results obtained, the following recommendations are forwarded: Biological yield of sugarcane for seed production required more NPSB fertilizer than what is being applied to attain its maximum. Hence, to get higher sett yield and ultimately seed cane of average quality it is recommended that application of 380 kg ha^-1^ of NPSB and 160 kg ha^-1^ Urea should be applied to seed cane plants of sugarcane variety NCO-334 and N-14. Whereas application of 320 kg ha^-1^ NPSB and 160 kg ha^-1^ urea for variety C86/56 is recommended under the study area. Hence, as this study is the first by its kind under Tana Beles condition,

**Figure 1:**
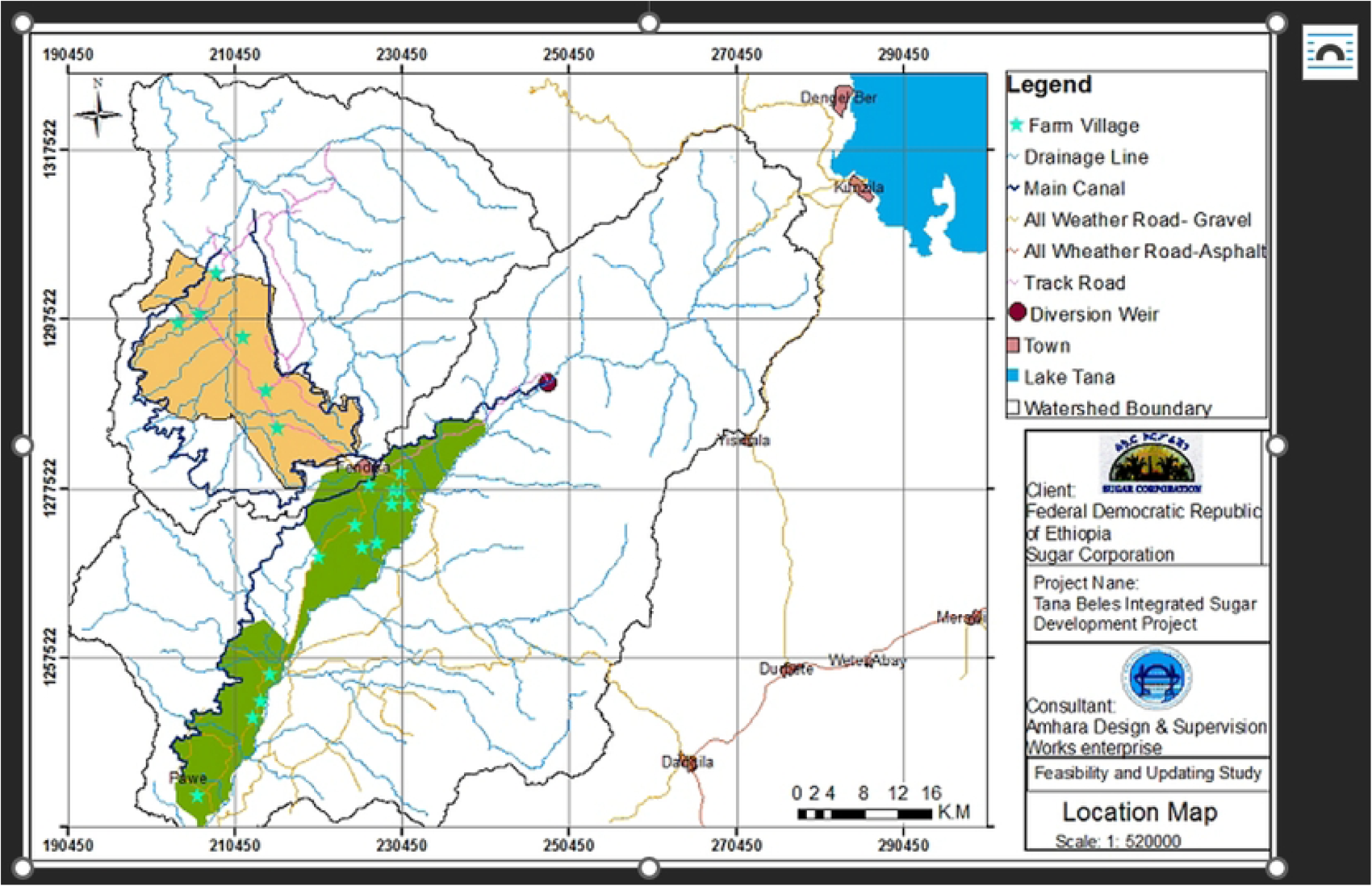
Location map of Tana Beles sugar development project

